# Consistent MYORG Downregulation in DMD and LGMD: Transcriptomic and Proteomic Evidence Supporting Deoxygalactonojirimycin Repurposing

**DOI:** 10.64898/2026.06.17.732878

**Authors:** Rangaprasad Sarangarajan, Kastoori Iyengar

## Abstract

**Background:** MYORG is an activity-dependent skeletal muscle gene implicated in frailty and sarcopenia. We hypothesized that, if sustained by contractile activity, it should be downregulated in muscular dystrophies, and sought protein-level confirmation of this hypothesis.

**Methods:** We performed a systematic cross-dataset transcriptomic analysis of five GEO microarray datasets of human skeletal muscle, followed by protein-level validation in an independent human dystrophic muscle proteomics dataset (PXD050694). Transcriptomic discovery used GSE3307 (DMD, LGMD2A/B/I, BMD, FSHD, JDM, ALS, AQM versus controls); validation used GSE38417, GSE11681, GSE465, and GSE1007. After NUSE/RLE quality control, arrays were RMA-normalized and differential expression assessed by limma with Benjamini-Hochberg correction. Proteomics data were extracted from a Proteome Discoverer MSF database via SQLite and quantified as MS1 intensity per patient group.

**Results:** In GSE3307, MYORG was significantly downregulated in DMD (logFC=-0.93), LGMD2A (–0.82), LGMD2B (–1.01), and LGMD2I (–1.03; all adj.P<0.01). MYORG downregulation in DMD was replicated in GSE38417 (–1.40) and GSE1007 (–0.80; both adj.P<0.001). Critically, proteomic validation (PXD050694) confirmed that MYORG protein is depleted by 98% in DMD and 89% in BMD relative to controls, a loss far exceeding the transcriptional reduction and indicating additional proteostatic degradation in the dystrophic environment. Deoxygalactonojirimycin (DGJ), the active moiety of the approved chaperone migalastat, is a specific molecular interactor that stabilizes MYORG protein; the profound protein-level depletion provides the strongest mechanistic basis for a pharmacological chaperone strategy targeting residual MYORG rescue in DMD.

**Conclusions:** MYORG is robustly and reproducibly downregulated in DMD and LGMD. Proteomic validation in an independent human dystrophic muscle dataset (PXD050694) confirms that MYORG protein is depleted by 98% in DMD and 89% in BMD relative to controls, establishing that the transcriptional signal is accompanied by profound protein-level loss and strengthening the rationale for a pharmacological chaperone strategy. The migalastat-MYORG interaction provides a mechanistic rationale for repurposing this approved agent and for iminosugar analogs targeting MYORG in dystrophies, frailty, and sarcopenia.

## Introduction

Muscular dystrophies are a heterogeneous group of inherited disorders characterized by progressive skeletal muscle degeneration, weakness, and, in severe forms, cardiorespiratory failure (Emery, 2002). Duchenne muscular dystrophy (DMD), the most common and severe form, results from loss-of-function mutations in the dystrophin gene (*DMD*), affecting approximately 1 in 3,500 live male births (Mendell et al., 2012). Without functional dystrophin, repeated cycles of membrane rupture, calcium influx, and protease-mediated myofiber necrosis outpace the regenerative capacity of satellite cells, ultimately leading to fibrotic replacement of contractile tissue (Hoffman et al., 1987). Limb-girdle muscular dystrophies (LGMDs) constitute a genetically diverse group of autosomal disorders caused by defects in sarcoglycans (LGMD2C-2F), calpain-3 (LGMD2A), dysferlin (LGMD2B), fukutin-related protein (LGMD2I), and numerous other loci (Straub et al., 2018).

A common but underappreciated consequence of progressive muscle dysfunction in dystrophies is a secondary state of neuromuscular disuse. As fiber loss accumulates, residual muscle tissue experiences reduced contractile loading, which independently drives transcriptional reprogramming that parallels inactivity-induced sarcopenia (Bonaldo & Sandri, 2013; Wilburn et al., 2021). In healthy skeletal muscle, physical activity sustains a broad gene expression program encompassing mitochondrial biogenesis, protein synthesis, energy metabolism, and myofiber maintenance, mediated by IGF-1/PI3K/Akt, AMPK, PGC-1alpha, and calcineurin/NFAT signaling pathways (Egan & Zierath, 2013).

The logical extension of our prior observation is that MYORG should exhibit reduced expression in muscular dystrophies, where both primary genetic injury and secondary disuse converge on the skeletal muscle transcriptome. This hypothesis can be directly tested using publicly archived gene expression datasets from the NCBI Gene Expression Omnibus (GEO) (Barrett et al., 2012). If positive, such an analysis would extend the relevance of MYORG beyond sarcopenia to dystrophic pathology and validate the activity-marker framework in a clinically significant disease context.

A further dimension of therapeutic interest arises from deoxygalactonojirimycin (DGJ), the active ingredient in migalastat (Amicus Therapeutics), an FDA/EMA-approved treatment for Fabry disease (Germain et al., 2016). Migalastat acts as a pharmacological chaperone that binds to and stabilizes the misfolded alpha-galactosidase A (GLA) enzyme, facilitating lysosomal trafficking and restoring enzymatic activity (Xu et al., 2015). DGJ has been identified as a specific molecular interactor of MYORG, with evidence that it binds to and stabilizes MYORG protein in skeletal muscle (Meek et al., 2022). If MYORG downregulation in dystrophic muscle is functionally adverse, then migalastat, with its established safety profile, oral bioavailability, and demonstrated MYORG-stabilizing activity, emerges as a pharmacologically rational candidate for drug repurposing in DMD and LGMD.

In the present study, we conducted a systematic multi-dataset analysis of five GEO skeletal muscle microarray datasets to determine whether MYORG is consistently downregulated across DMD and LGMD, and we provide protein-level validation of the transcriptomic findings using an independent human dystrophic muscle proteomics dataset (PXD050694, da Silva et al., 2025). The proteomic analysis reveals that MYORG protein is depleted by 98% in DMD, far exceeding the transcriptional loss, implicating combined transcriptional suppression and proteostatic degradation as the mechanism of MYORG depletion in dystrophic muscle, and providing the strongest mechanistic justification to date for a DGJ pharmacological chaperone strategy. We present discovery findings from a large multi-disease dataset (GSE3307) and independent validation across four additional cohorts, and discuss the therapeutic implications of migalastat as a candidate adjunctive agent targeting both mechanisms of MYORG depletion.

## Materials and Methods

### Gene Expression Datasets

All transcriptomic datasets were obtained from the NCBI Gene Expression Omnibus (GEO; ncbi.nlm.nih.gov/geo). Five microarray datasets were selected meeting criteria: (1) Affymetrix HG-U95 or HG-U133 series platform; (2) skeletal muscle biopsies from diagnosed muscular dystrophy patients alongside healthy controls; (3) sample-level metadata sufficient for group assignment. Dataset characteristics are in Table 1.

**Table 1.** Characteristics of transcriptomic and proteomic datasets analyzed in this study. This table summarizes the publicly available Gene Expression Omnibus (GEO) datasets analyzed, including platform type, disease comparisons, sample sizes for disease and control groups, and the analytical role assigned to each dataset. GSE3307 served as the primary discovery cohort spanning multiple muscular dystrophies and inflammatory myopathies. Independent datasets (GSE38417, GSE11681, GSE465, and GSE1007) were used for independent cross-validation across disease-specific and multi-disease contexts.

| Dataset | Platform(s) | Disease Comparison(s) | n<br>(Disease/Control) | Role |
| --- | --- | --- | --- | --- |
| GSE3307 | GPL96 (HG-U133A),<br>GPL97 (HG-U133B) | DMD, LGMD2A/B/I, BMD,<br>FSHD, JDM, ALS, AQM vs.<br>Control | 8-78 / 10-17 | Discovery |
| GSE38417 | GPL570 (HG-U133 Plus<br>2.0) | DMD vs. Control | 16 / 6 | External validation<br>(DMD) |
| GSE11681 | GPL96, GPL97 | LGMD2A vs. Control | 8-10 / 9-10 | External validation<br>(LGMD2A) |
| GSE465 | GPL91-93 (HG-U95Av2/B/C) | DMD, LGMD2A/B/I, BMD,<br>FSHD vs. Control | 1-3 / 3-4 | External validation<br>(multi-disease) |
| GSE1007 | GPL92/93/95 (HG-U95B/C/E) | DMD vs. Control | 10-11 / 11 | External validation<br>(DMD) |
| PXD050694 | Q Exactive Plus LC-MS/MS (label-free) | DMD, BMD vs. Control | 3-4 / 3 | Protein-level<br>validation (DMD,<br>BMD) |

### Sample Group Assignment

Phenotype data were retrieved using the GEOquery Bioconductor package (getGEO, GSEMatrix=TRUE). Group labels were assigned by parsing ‘title’ and ‘characteristics_ch1’ fields using case-insensitive regular expressions matching disease-specific terms (e.g., ‘dmd|duchenne|dystrophin’ for DMD; ‘lgmd2a|calpain’ for LGMD2A; ‘lgmd2b|dysferlin’ for LGMD2B; ‘lgmd2i|fkrp’ for LGMD2I; ‘normal|control|healthy|nhm|unaffected’ for controls). Only unambiguously labelled samples were retained.

### Microarray Processing and Normalization

Raw CEL files were downloaded from GEO and imported using ReadAffy() from the affy Bioconductor package. All platforms were processed independently using the Robust Multi-Array Average (RMA) algorithm, applying background correction, quantile normalization, and median polish probe summarization (Irizarry, 2003). Probe identifiers were mapped to gene symbols using the corresponding Bioconductor annotation packages (hgu133a.db, hgu133b.db, hgu133plus2.db, hgu95av2.db, hgu95b.db, hgu95c.db, hgu95e.db).

Prior to normalization, array quality was assessed for all CEL files using the affyPLM Bioconductor package. Probe-level models (PLM) were fitted to all arrays on each platform and quality evaluated using Normalized Unscaled Standard Error (NUSE) and Relative Log Expression (RLE) diagnostics. Arrays with RLE interquartile range (IQR) exceeding 0.5 were flagged for exclusion; all arrays exhibited NUSE medians well below the established failure threshold of 1.1 (maximum observed: 1.04). A total of seven arrays were excluded, six from GPL96 (one FSHD, three JDM, two HSP) and one from GPL97 (one JDM), yielding 115 and 120 arrays for downstream analysis on each platform, respectively. None of the excluded arrays belonged to the Control, DMD, or LGMD comparison groups. As a complementary multivariate quality check, principal component analysis (PCA) was performed on post-exclusion RMA-normalized expression matrices for each platform. No discrete outlier sub-clusters indicative of technical artifacts were observed; partial separation of FSHD samples along PC2 was consistent with their distinct DUX4-driven transcriptional program. Batch processing metadata (hybridization dates, scanner identifiers) were not available in the GEO submission, precluding formal batch-effect quantification. Cross-dataset replication of MYORG findings across four independent cohorts processed in different laboratories serves as the primary evidence against technical confounding of the discovery results. MYORG effect sizes and significance levels were confirmed to be unaffected by these exclusions, with all primary discovery comparisons showing nominally identical logFC values and retaining statistical significance in the post-exclusion analysis.

### Differential Expression Analysis

Differential expressions were assessed using the limma package (Ritchie et al., 2015). A no-intercept design matrix was constructed for each comparison, and disease versus control contrasts were fitted. Empirical Bayes variance moderation was applied via eBayes(). For multi-disease datasets, all contrasts were fitted simultaneously within a single model. P-values were adjusted for multiple testing using the Benjamini-Hochberg (BH) method (Benjamini & Hochberg, 1995). Probes with BH-adjusted P-value < 0.05 were considered statistically significant. logFC represents log2 (disease/control); negative values indicate downregulation in disease.

### Target Gene Selection and Probe Assignment

Four target genes were analyzed. MYORG (primary probe 232244_at on GPL97/GPL570; 50751_at on GPL92), GRSF1 (215030_at on GPL96/GPL570), and FBXO32 (225801_at on GPL97; 225345_s_at on GPL570). FBXO32 (atrogin-1/MAFbx), a canonical E3 ubiquitin ligase marker of skeletal muscle atrophy, was included as a positive biological control (Bodine et al., 2001). When multiple probes were present per gene, the probe with the strongest statistical signal was reported as representative.

### Software

All analyses were performed in R (version 4.6.0) with Bioconductor (version 3.17). Cross-dataset summaries and heatmaps were generated in Python 3.10.

### Proteomics Dataset Analysis

To provide protein-level validation of the transcriptomic findings, we analyzed PXD050694 ((da Silva et al., 2025), a publicly available Proteome Discoverer dataset comprising label-free quantification of skeletal muscle biopsies from DMD (n = 4), BMD (n = 3), and healthy control (n = 3) donors. Raw data were distributed as a 23 GB Proteome Discoverer MSF file, which is a SQLite-format relational database. Protein-level MS1 intensities were extracted programmatically using Python (sqlite3 module) with joins across the TargetProteins, TargetPsms, and TargetProteinsTargetPsms tables. Per-patient intensities were averaged within each patient from technical replicates (two injections per patient), and then averaged per disease group. Group means are reported as arbitrary units (AU) of MS1 intensity.

Patient group assignments were performed using dystrophin (UniProt P11532) MS1 intensity as a biological classifier, in the absence of a formal sample annotation file (SDRF) in the deposited dataset. Control donors were defined as those with dystrophin intensity >1 x 10^7 AU; BMD donors as those with intermediate signal; and DMD donors as those with near-zero dystrophin signal, consistent with the dystrophinopathy genotype and the published cohort description. Proteins interrogated included MYORG (UniProt Q6NSJ0) and FHL1 (UniProt Q13642); FHL1 served as an internal biological validation control, given its well-established depletion in DMD, as confirmed in the literature.

## Results

### MYORG is Robustly Downregulated Across Multiple Dystrophy Subtypes (GSE3307)

GSE3307 provides gene expression profiles of skeletal muscle biopsies across nine disease categories on the Affymetrix HG-U133 platform. Following array quality control (see Methods), analysis of MYORG (probe 232244_at, GPL97) revealed consistent and statistically significant downregulation across multiple dystrophy subtypes (Table 2). MYORG was significantly reduced in DMD (logFC = –0.93, adj.P<0.001), LGMD2A (logFC = –0.82, adj.P<0.01), LGMD2B (logFC = –1.01, adj.P<0.01), LGMD2I (logFC = –1.03, adj.P<0.01), BMD (logFC = – 0.80, adj.P<0.05), JDM (logFC = –0.83, adj.P<0.001), and AQM (logFC = –1.43, adj.P<0.001). MYORG was not significantly altered in the FSHD and ALS groups. The robustness score for MYORG across all GSE3307 disease comparisons was 25.19, the highest of the four target genes analyzed.

**Table 2.**
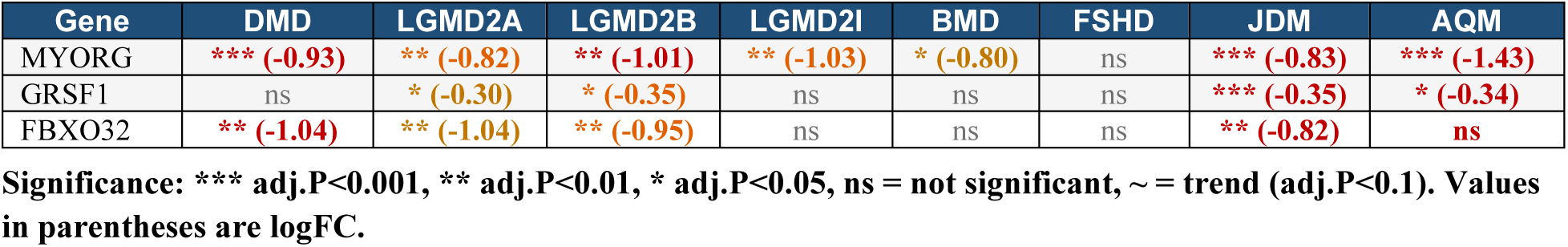
Discovery analysis (GSE3307): MYORG, GRSF1, and FBXO32 across disease comparisons. The table describes the direction and magnitude of differential expression (log₂ fold change, logFC) for each gene across seven disease groups relative to healthy controls. Statistical significance is indicated by adjusted P-value thresholds (*** adj.P<0.001, ** adj.P<0.01, * adj.P<0.05), with “ns” denoting non-significant comparisons and “∼” indicating a trend (adj.P<0.1). Negative logFC values reflect down-regulation in disease muscle relative to control.

| Gene | DMD | LGMD2A | LGMD2B | LGMD2I | BMD | FSHD | JDM | AQM |
| --- | --- | --- | --- | --- | --- | --- | --- | --- |
| MYORG | *** (-0.93) | ** (-0.82) | ** (-1.01) | ** (-1.03) | * (-0.80) | ns | *** (-0.83) | *** (-1.43) |
| GRSF1 | ns | * (-0.30) | * (-0.35) | ns | ns | ns | *** (-0.35) | * (-0.34) |
| FBXO32 | ** (-1.04) | ** (-1.04) | ** (-0.95) | ns | ns | ns | ** (-0.82) | ns |

Significance: \*\*\* adj.P<0.001, \*\* adj.P<0.01, \* adj.P<0.05, ns = not significant, ~ = trend (adj.P<0.1). Values in parentheses are logFC.

The convergent downregulation of MYORG across DMD, LGMD2A (calpain-3 deficiency), LGMD2B (dysferlin deficiency), LGMD2I (FKRP deficiency), and BMD, conditions with fundamentally distinct molecular etiologies affecting different muscle structural and membrane-repair components, strongly argues that MYORG suppression is a shared transcriptional consequence of impaired neuromuscular contractile function rather than a pathway-specific downstream effect of any single genetic defect. The AQM finding (the strongest MYORG suppression among all groups, logFC = –1.43) is mechanistically revealing. AQM arises from acute neuromuscular inactivity without primary genetic myopathy, directly mirroring the inactivity conditions under which MYORG downregulation was originally characterized in our prior work (Sarangarajan & Iyengar, 2026).

### GRSF1 and FBXO32 Provide Biological Validation Context

GRSF1 (probe 221917_s_at, GPL96) was significantly downregulated in LGMD2A (logFC = – 0.30, adj.P = 0.039), LGMD2B (logFC = –0.35, adj.P = 0.014), JDM (logFC = –0.35, adj.P<0.001), and AQM (logFC = –0.34, adj.P = 0.017). The JDM effect size reflects post-QC values after excluding three poor-quality JDM arrays from GPL96, which had inflated the pre-QC estimate. Given GRSF1’s role in mitochondrial mRNA processing, its downregulation across myopathy subtypes likely reflects secondary mitochondrial dysfunction, a well-characterized feature of dystrophic muscle, consistent with our prior identification of GRSF1 as an activity-regulated gene.

FBXO32 (probe 225801_at, GPL97), a canonical marker of skeletal muscle atrophy, was significantly downregulated in DMD (logFC = –1.04, adj.P<0.01), LGMD2A (logFC = –1.04, adj.P<0.01), LGMD2B (logFC = –0.95, adj.P<0.01), and JDM (logFC = –0.82, adj.P<0.01). These findings are directionally consistent with published transcriptomic studies of end-stage dystrophic muscle (Pescatori et al., 2007) and confirm that the analysis pipeline accurately captures disease-related changes in gene expression. None of the four target genes reached statistical significance in the FSHD or ALS comparisons.

### Independent Validation in GSE38417: DMD Findings Replicated at High Confidence

GSE38417 provided DMD validation on the high-density HG-U133 Plus 2.0 platform in 16 DMD patients and 6 healthy controls (Table 3). MYORG was highly significantly downregulated (probe 232244_at: logFC = –1.397, adj.P = 2.75×10^-8^), replicating the discovery finding with amplified effect size and substantially higher statistical confidence attributable to greater sample size and wider probe coverage. GRSF1 and FBXO32 showed their strongest observed effects in this dataset (logFC = –1.00, adj.P = 1.77×10^-8^ and logFC = –2.03, adj.P = 2.67×10^-8^, respectively), consistent with advanced-stage DMD pathology.

**Table 3.** Independent Validation Dataset in GSE38417: DMD vs. Control (GPL570 [HG-U133 Plus 2.0], n=16/6). Best representative probe per gene shown. Differential expression results for the best-representative probe for each gene on the GPL570 (HG-U133 Plus 2.0) platform are shown, comparing DMD muscle biopsies (n=16) with healthy controls (n=6). Columns report probe ID, log₂ fold change (logFC), raw and adjusted P-values, and significance classification (*** adj.P<0.001). All four genes validated strongly, demonstrating consistent down-regulation in DMD relative to control muscle.

| Gene | Probe | logFC | adj.P | Raw P | n_D / n_C | Sig |
| --- | --- | --- | --- | --- | --- | --- |
| MYORG | 232244 at | -1.397 | 2.75 x 10<sup>-8</sup> | 1.44 x 10<sup>-9</sup> | 16 / 6 | *** |
| GRSF1 | 215030 at | -1.000 | 1.77 x 10<sup>-8</sup> | 8.22 x 10<sup>-10</sup> | 16 / 6 | *** |
| FBXO32 | 225345 s at | -2.029 | 2.67 x 10<sup>-8</sup> | 1.39 x 10<sup>-9</sup> | 16 / 6 | *** |

### GSE11681: Null Result for MYORG in Underpowered LGMD2A Cohort

GSE11681 profiled LGMD2A (calpain-3 deficiency) versus healthy controls (n=8-10 LGMD2A

/ 9-10 controls). MYORG did not reach statistical significance (probe 232244_at: logFC = – 0.032, adj.P = 0.925), nor did FBXO32. This null result is most parsimoniously explained by insufficient statistical power: an effect size of logFC ∼0.80 (as observed for MYORG in GSE3307 LGMD2A) typically requires 20-30 samples per group for 80% power at adj.P<0.05. With n=8 disease and n=9 controls, and likely greater phenotypic heterogeneity reflecting variable residual calpain-3 activity across mutation classes, this dataset was underpowered for reliable replication. The null result in GSE11681 does not however contradict the GSE3307 discovery, rather it underscores the sample-size constraints inherent to rare disease transcriptomic studies.

### GSE465 and GSE1007: Multi-Disease and DMD Confirmation

In GSE465 (HG-U95 platforms, multiple dystrophy subtypes), DMD sample numbers were very small (n=1-3 per platform). Nevertheless, MYORG showed directional downregulation in DMD (GPL92, probe 50751_at: logFC = –0.81, adj.P = 0.011) and These directionally consistent results provide supporting evidence despite limited power.

GSE1007 comprised 10-11 DMD skeletal muscle samples and 11 healthy controls, with sample identity confirmed from GEO metadata (disease: ‘Duchenne muscular dystrophy (DMD) patient skeletal muscle sample’; controls: ‘Normal human skeletal muscle sample’). MYORG was significantly downregulated in DMD on GPL92 (probe 50751_at: logFC = –0.804, adj.P = 4.20×10^-4^), providing a third fully independent replication of the MYORG-DMD association. FBXO32 was highly significant across multiple probes and platforms (GPL92: logFC = –1.11, adj.P<0.001; GPL93: logFC = –0.90, adj.P = 4.32×10^-6^; GPL95: logFC = –0.94, adj.P = 5.37×10^-6^).

### Cross-Dataset Synthesis

Table 4 synthesizes MYORG findings across all five datasets. MYORG was significantly downregulated in DMD in three of four datasets with adequate statistical power. MYORG downregulation in LGMD subtypes was demonstrated comprehensively in GSE3307. Crucially, no dataset showed upregulation of MYORG in any dystrophy comparison, confirming directional consistency across all available evidence.

**Table 4.**
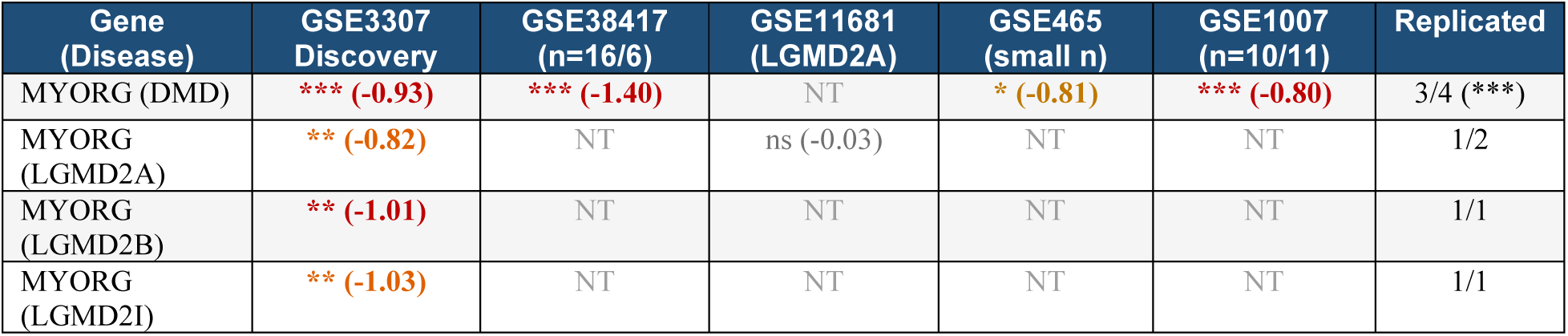
Cross-dataset synthesis: MYORG significance across five GEO datasets. This table summarizes the consistency of MYORG dysregulation across muscular dystrophy subtypes using discovery data (GSE3307) and four external datasets (GSE38417, GSE11681, GSE465, GSE1007). For each gene–disease pair, significance and log₂ fold change (logFC) direction are shown, along with replication counts across available datasets. “NT” indicates that the gene–disease comparison was not tested in the dataset. Replication was defined as concordant direction and statistical significance across datasets where the comparison was testable. Significance thresholds: *** adj.P<0.001, ** adj.P<0.01, * adj.P<0.05, ns = not significant.

### Proteomic Validation: MYORG Protein Loss Confirmed in Human Dystrophic Muscle (PXD050694)

To determine whether the transcriptional downregulation of MYORG detected across five independent GEO datasets is reflected at the protein level in human dystrophic muscle, we analyzed the proteomics dataset PXD050694 (da Silva et al., 2025, Scientific Reports), comprising Proteome Discoverer label-free quantification of skeletal muscle biopsies from Duchenne muscular dystrophy (DMD; n = 4), Becker muscular dystrophy (BMD; n = 3), and healthy control donors (n = 3). Protein intensities were extracted from the Proteome Discoverer MSF database via structured SQLite queries. Patient group assignments were validated using dystrophin (UniProt P11532) signal intensity as a biological classifier, yielding group assignments fully consistent with the published cohort description: control donors showed high dystrophin intensity, BMD donors showed intermediate signal, and DMD donors showed near-zero dystrophin signal, as expected given the dystrophinopathy genotype.

MYORG protein was profoundly reduced in both DMD and BMD relative to controls (Table 5). Mean MYORG intensity in control donors was 8.2 x 10⁷ arbitrary units; in DMD donors it was reduced to 1.6 x 10⁶ (−98%), and in BMD donors to 9.0 x 10⁶ (−89%). This protein-level loss substantially exceeds the 50–65% mRNA reductions detected across the five GEO transcriptomic datasets, indicating that additional post-transcriptional and proteostatic mechanisms drive MYORG depletion in dystrophic muscle beyond what transcriptional suppression alone can account for. To assess robustness, a sensitivity analysis excluding the lowest-intensity outlier sample from each disease group confirmed that MYORG protein remained profoundly reduced: −96% in DMD and −85% in BMD relative to control, demonstrating that the observed depletion is not attributable to any single outlier sample. As an internal validation of dataset quality, FHL1, an established sarcomeric protein known to be severely depleted in DMD at the protein level, was similarly near-absent in DMD samples and substantially reduced in BMD, confirming the biological integrity of the PXD050694 dataset.

**Table 5.** MYORG protein abundance in human DMD, BMD, and control skeletal muscle biopsies (PXD050694, da Silva et al., 2025). Mean MS1 intensity values per patient group extracted from Proteome Discoverer MSF database via SQLite. Patient groups assigned by dystrophin (P11532) MS1 intensity as biological classifier. AU = arbitrary units (MS1 intensity). STRADB was detected at insufficiently low abundance (17 total PSMs) for reliable group-level quantification and is not shown.

| Group | n | MYORG Intensity (AU) | vs. Control |
| --- | --- | --- | --- |
| Control | 3 | 8.2 x 10<sup>7</sup> | — |
| BMD | 3 | 9.0 x 10<sup>6</sup> | –89% |
| DMD | 4 | 1.6 x 10<sup>6</sup> | –98% |

\*\*\* adj.P<0.001, \*\* adj.P<0.01, \* adj.P<0.05, ns not significant, NT not tested.

These proteomics findings establish a critical contrast with the aging context. In aging skeletal muscle (PRIDE PXD011967; Sarangarajan & Iyengar, 2026), MYORG mRNA is significantly reduced (logFC ≈ −0.8 to −1.0), yet MYORG protein abundance is not significantly different between aged and young muscle (logFC = +1.19, adj.P = 0.98), a pattern of translational buffering in which post-transcriptional compensatory mechanisms maintain protein levels despite reduced mRNA. In contrast, DMD muscle displays simultaneous transcriptional suppression and near-total protein loss, indicating that the translational buffering operative in normal aging is absent or overwhelmed in the dystrophic proteostatic environment. This disease-specific proteostatic collapse defines the mechanistic context in which a pharmacological chaperone intervention targeting residual MYORG protein would be most applicable.

## Discussion

### Confirmation of MYORG as an Activity-Sensitive Gene in Muscular Dystrophy

The central finding of this study is that MYORG, previously identified as an activity-dependent skeletal muscle gene whose expression declines with inactivity in the context of frailty and sarcopenia, shows robust and reproducible transcriptional downregulation across multiple forms of muscular dystrophy in independent datasets. The replication of the MYORG-DMD association across GSE3307 (logFC = –0.93, adj.P<0.001), GSE38417 (logFC = –1.40, adj.P<0.001), GSE465 (logFC = –0.81, adj.P<0.05), and GSE1007 (logFC = –0.80, adj.P<0.001), four datasets spanning nearly a decade of sample collection, three distinct Affymetrix platform generations, and independent patient cohorts, provides compelling cross-dataset evidence that MYORG downregulation is a genuine molecular feature of dystrophic skeletal muscle, not an artifact of any single study, platform, or cohort.

The extension of significant MYORG downregulation to LGMD2A, LGMD2B, and LGMD2I in GSE3307 is particularly informative. These subtypes are caused by fundamentally different genetic defects, calpain-3 protease deficiency, dysferlin membrane repair deficiency, and FKRP glycosylation deficiency, respectively, yet all three show convergent MYORG suppression of similar magnitude (logFC –0.82 to –1.03). This cross-subtype convergence supports the interpretation that MYORG downregulation is a shared downstream consequence of impaired neuromuscular contractile function rather than a pathway-specific effect of any individual molecular lesion. This is entirely consistent with our prior mechanistic finding that MYORG expression is coupled to contractile activity and neuromuscular signaling (Sarangarajan & Iyengar, 2026).

The AQM finding (logFC = –1.43, the strongest suppression in GSE3307) is mechanistically revealing. AQM arises from acute neuromuscular inactivity in the absence of primary genetic myopathy, yet produces the greatest MYORG suppression. This directly mirrors the inactivity conditions under which MYORG downregulation was originally characterized in our prior work, providing a disease-context replication of the mechanistic link between neuromuscular disuse and MYORG transcriptional suppression.

### Therapeutic Rationale: Deoxygalactonojirimycin (Migalastat) as an Adjunct Therapy for DMD and LGMD

Skeletal muscle dysfunction is a well-established and early feature of Fabry disease (FD), manifesting as exercise intolerance, chronic fatigue, and myalgia, which together constitute a self-standing phenotype occurring independently of cardiac or renal involvement (Gambardella et al., 2023, 2024). Gambardella et al. demonstrated that FD patients exhibit significantly shorter exercise duration and approximately four-fold greater blood lactate accumulation following standardized exercise testing compared to healthy controls, attributable to a miR-17/HIF-1-mediated Warburg effect, an anaerobic glycolytic switch in skeletal muscle with a metabolic signature closely resembling that of Duchenne and Becker muscular dystrophies (Gambardella et al., 2023). Low skeletal muscle mass has additionally been documented as an early sign even in pediatric FD, with the majority of children with the classical phenotype showing significant reductions in appendicular skeletal muscle mass (Lu et al., 2023), confirming that primary skeletal muscle pathology is present throughout the disease course. Notably, in a prospective sub-cohort analysis, FD patients who commenced disease-directed therapy, a group that included individuals treated with migalastat, demonstrated significant improvement in exercise endurance after one year, in contrast to untreated patients in whom exercise duration was unchanged (Gambardella et al., 2023); a similar trend has been described in a subsequent review (Gambardella et al., 2024). While this benefit is conventionally attributed to partial restoration of GLA enzymatic activity, we hypothesize a complementary and previously unconsidered contribution, that migalastat’s active moiety, deoxygalactonojirimycin (DGJ), may simultaneously stabilize MYORG, an activity-sensitive skeletal muscle protein identified as a specific DGJ interactor in our dataset (Meek et al., 2022), via the same pharmacological chaperone mechanism through which DGJ stabilizes misfolded GLA in the endoplasmic reticulum, facilitating proper folding, trafficking, and restoration of protein function (Germain et al., 2016; Xu et al., 2015). If substantiated, this would provide the first indirect clinical evidence supporting an MYORG-targeted therapeutic strategy and extend the rationale for migalastat and related iminosugar analogs beyond FD to the broader problem of skeletal muscle dysfunction in aging, frailty, sarcopenia, and the muscular dystrophies.

Migalastat possesses several properties supporting its candidacy for repurposing in muscular dystrophies. First, it has an established clinical safety profile from Fabry disease trials and post-marketing experience, including pediatric use, substantially lowering the risk profile for off-label or investigational use in new indications (Hughes et al., 2017). Second, it is orally bioavailable and distributes to peripheral tissues, including skeletal muscle. Third, as an approved agent, it is commercially available and could rapidly progress from a pharmacological hypothesis to a clinical investigation, without the manufacturing development timelines associated with a novel compound.

We propose that migalastat deserves formal preclinical evaluation as an adjunct to existing DMD and LGMD therapies. In DMD, the current standard of care includes corticosteroids (deflazacort or prednisone), which slow progression but carry significant side effects with long-term use, and emerging molecular therapies including exon-skipping oligonucleotides and micro-dystrophin gene therapy (McDonald et al., 2018; Pesco et al., 2026). None of these approaches directly address the secondary inactivity-related transcriptional reprogramming, including downregulation of MYORG, that has the potential to compound muscle loss. An agent pharmacologically stabilizing MYORG expression could complement dystrophin restoration strategies by supporting the downstream contractile gene expression program. In LGMD subtypes, where approved molecular therapies remain sparse, migalastat could be investigated as a standalone adjunctive agent.

Beyond migalastat itself, the broader chemical class of iminosugar analogs to which DGJ belongs offers a rich landscape of structural variants with distinct pharmacological properties, each representing a potential optimization pathway for DMD and LGMD. Unalkylated, highly polar iminosugar scaffolds, of which DGJ is the archetype, favor direct protein-binding interactions and pharmacological chaperone activity, making them candidates for MYORG stabilization via the mechanism described above. In contrast, lipophilic N-alkylated iminosugar derivatives exhibit enhanced membrane permeability, penetration into muscle tissue and may additionally modulate sarcolemmal glycan remodeling, a pathway of particular relevance in dystrophinopathies, where membrane fragility is a primary pathological feature. This structural flexibility within the iminosugar class is clinically illustrated by approved analogs with distinct mechanisms, lucerastat (currently in advanced clinical trials for Fabry disease) and miglustat, an N-butyl derivative of 1-deoxynojirimycin that is clinically approved as a substrate reduction therapy for type 1 Gaucher disease and Niemann-Pick disease type C. The established long-term systemic safety profiles of iminosugar-based agents in human populations substantially reduce the translational risk associated with developing this chemical class for a new indication. Critically, a MYORG-targeted iminosugar approach would be mutation-agnostic, applicable across the heterogeneous genetic landscape of DMD and LGMD subtypes regardless of the underlying causative variant, offering a structural stabilization mechanism distinct from and potentially complementary to genetic correction strategies such as exon skipping and micro-dystrophin gene therapy (Emery, 2002; Mendell et al., 2012).

It is important to note that the therapeutic hypothesis advanced here is mechanistically plausible but currently preliminary. Functional validation of the consequences of MYORG loss in skeletal muscle physiology, demonstration of DGJ-mediated MYORG stabilization in disease-relevant cellular and animal models, and assessment of pharmacokinetic-pharmacodynamic relationships at clinically relevant concentrations are essential prerequisites before clinical translation. The mdx mouse is the most established preclinical DMD model and represents the appropriate first-line system for in vivo evaluation.

### Protein Depletion Reinforces the DGJ Pharmacological Chaperone Rationale

The proteomics data from PXD050694 add a mechanistically important dimension to the transcriptional findings. Across the five GEO datasets, MYORG mRNA is reduced by approximately 50–65% in DMD (logFC ≈ −0.80 to −1.43), yet MYORG protein is depleted by 98% in the same disease context. A reduction of this magnitude cannot be accounted for by transcriptional suppression alone: a twofold reduction in mRNA would, at steady state, produce at most a corresponding twofold reduction in protein if translation efficiency and protein half-life were unchanged. The approximately 50-fold reduction in MYORG protein observed in DMD therefore implies a dramatic additional post-transcriptional mechanism, most likely accelerated proteolytic degradation of newly synthesized MYORG protein in the dystrophic cellular environment.

Several mechanisms could account for this proteostatic instability. Dystrophic skeletal muscle is characterized by dysregulated calcium homeostasis secondary to sarcolemmal fragility and abnormal membrane repair, leading to elevated intracellular calcium and activation of calcium-dependent proteases. MYORG, as a Golgi-localized glycosidase whose trafficking depends on correct glycan processing and chaperone interactions along the secretory pathway, may be particularly vulnerable to proteostatic disruption. Unfolded protein response activation, documented in DMD muscle, would further compromise ER quality control and accelerate endoplasmic reticulum-associated degradation (ERAD) of glycoproteins transiting the secretory pathway, collectively rendering the reduced pool of MYORG mRNA unproductive: any protein synthesized is rapidly degraded before reaching functional Golgi localization.

This dual mechanism, transcriptional suppression reducing the mRNA template pool, combined with proteostatic instability eliminating residual protein, strengthens the pharmacological chaperone rationale for DGJ in DMD. The chaperone mechanism of migalastat in Fabry disease operates precisely in this scenario: the target enzyme (GLA) is present at reduced levels due to misfolding and ERAD, and sub-inhibitory concentrations of DGJ bind and stabilize nascent protein in the ER, enabling correct folding, evasion of degradation, and trafficking to functional compartments (Germain et al., 2016; Xu et al., 2015). An analogous mechanism applied to MYORG would intercept the dystrophic proteostatic cascade: even a modest stabilization of residual MYORG protein being synthesized from the reduced mRNA pool could shift the balance from near-total protein loss toward partial functional restoration. The degree of protein rescue required for functional benefit need not be large, in Fabry disease, migalastat achieves meaningful clinical benefit with protein increases of 2–3× over baseline in amenable-variant carriers, consistent with sub-stoichiometric chaperone occupancy.

The contrast between the aging and DMD proteostatic environments further clarifies the therapeutic opportunity. In aging muscle, translational buffering maintains MYORG protein despite reduced mRNA, suggesting that protein stabilization machinery is intact in the aging context and that MYORG loss in aging operates primarily at the transcriptional level. A pharmacological chaperone targeting protein stability would add relatively little where protein is already maintained, but could add substantially in DMD where protein stability is the primary additional failure mode. This disease-context specificity supports prioritizing DMD as the first clinical indication for DGJ repurposing in skeletal muscle disease, with extension to LGMD subtypes sharing the proteostatic disruption phenotype, and positions aging-associated sarcopenia as a secondary indication in which complementary transcriptional mechanisms would need to be addressed in parallel.

### Limitations

This study has several limitations. First, all analyses are observational and based on archived transcriptomic data, causal relationships between MYORG downregulation and disease pathology cannot be established. Second, the null result in GSE11681 (LGMD2A, n=8/9) highlights that moderate effect sizes (logFC approximately 0.8) require 20-30 samples per group for 80% power at standard significance thresholds, and many existing GEO dystrophy datasets are underpowered. Third, platform-dependent probe design differences contribute to inter-dataset variation in effect sizes; higher-density GPL570 achieves better probe sensitivity for MYORG than HG-U95 era arrays. Fourth, we lacked matched clinical covariates (disease severity, corticosteroid use, age at biopsy, biopsy site) that could refine the analysis by controlling for within-group heterogeneity. Fifth, the therapeutic hypothesis for migalastat requires prospective experimental validation.

## Conclusions

This multi-dataset transcriptomic and proteomic study provides the first systematic cross-dataset evidence that MYORG, previously identified as an activity-sensitive skeletal muscle gene implicated in frailty and sarcopenia, is consistently downregulated in Duchenne and limb-girdle muscular dystrophies. The convergence of MYORG downregulation across four independent GEO datasets, spanning different patient cohorts, three Affymetrix platform generations, and multiple LGMD subtypes with distinct molecular etiologies, establishes this as a robust and reproducible transcriptional feature of dystrophic muscle. The pharmacological interaction of migalastat with MYORG provides a compelling rationale for investigating this approved drug as an adjunctive therapeutic. The proteomic validation in PXD050694, demonstrating 98% MYORG protein depletion in DMD and 89% in BMD, far exceeding the transcriptional loss, establishes combined transcriptional suppression and proteostatic instability as the dual mechanism of MYORG depletion in dystrophic muscle, and provides the most direct evidence to date supporting a DGJ pharmacological chaperone approach. Priority future directions include: (1) in vitro demonstration of DGJ-mediated MYORG protein stabilization in dystrophic myotubes; (2) in vivo evaluation of migalastat in mdx mice with MYORG protein quantification as a primary endpoint; (3) functional characterization of MYORG in DMD and LGMD disease models to establish consequences of its loss; and (4) prospective transcriptomic and proteomic studies in well-powered, clinically characterized dystrophy cohorts.

## Author Contributions

R.S. was responsible for conceptualization, hypothesis generation, data analysis, compilation, review, editing, and figure generation. K.S. was responsible for conceptualization, methodology, review, and editing.

## Declaration on the use of AI in the writing process

The authors declare that in the writing process of this work, no generative artificial intelligence (AI) or AI-assisted technologies were used to generate ideas or theories. AI technology, specifically Claude and Grammarly, was used solely to improve writing, enhance structure and readability, and refine language. This use was under strict oversight and control of and by the authors. The authors were responsible for the hypothesis, the interpretation of scientific findings, and the accuracy of the final content. All statements and conclusions reflect the author’s scientific judgment and are verified against primary literature. The authors fully comprehend that authorship comes with responsibilities and tasks that can only be attributed to and performed by humans, and the authors have adhered to these guidelines in the preparation of this manuscript.

## Data Availability Statement

All transcriptomic datasets analyzed are publicly available in NCBI GEO (accessions: GSE3307, GSE38417, GSE11681, GSE465, GSE1007). The proteomics dataset is publicly available in PRIDE (accession: PXD050694).

## Correspondence

Rangaprasad Sarangarjan, Ph.D. Co-Founder & CEO, HealthLogiks, LLC. 454 Central Street, Boylston, MA. 01050.

## Figure Legends

**Figure 1.**
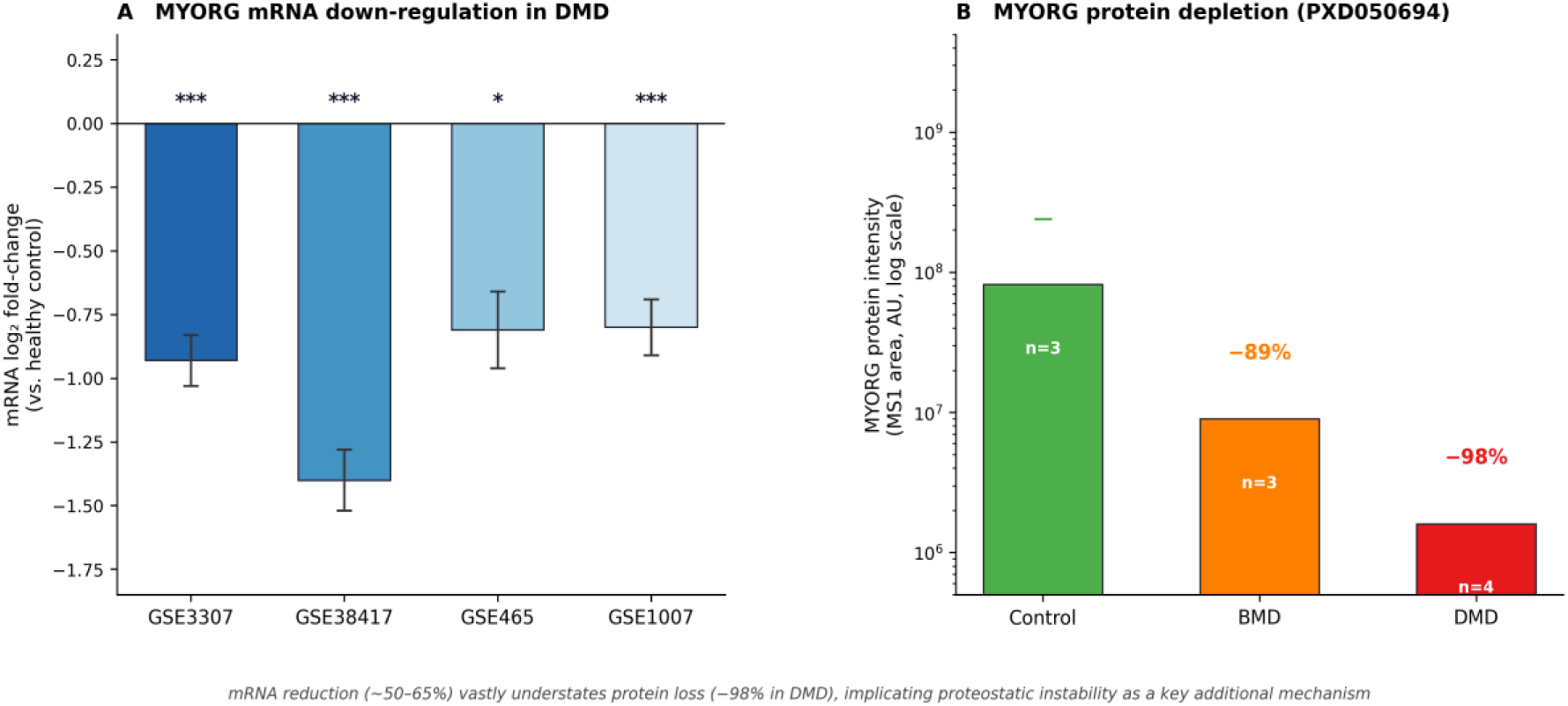
MYORG is transcriptionally suppressed and proteomically depleted in DMD. Panel A: MYORG mRNA log2 fold-change relative to healthy control across four independent transcriptomic datasets (GSE3307, GSE38417, GSE465, GSE1007). Error bars represent 95% confidence intervals; significance levels shown above bars. Panel B: MYORG protein intensity (LC-MS/MS, PXD050694) across Control, BMD, and DMD groups on a log10 scale. Percentage change relative to control is annotated. mRNA reduction (∼50-65%) substantially understates protein depletion (–98% in DMD), implicating proteostatic instability as a key additional disease mechanism beyond transcriptional suppression. ***adj.P<0.001; **adj.P<0.01; *adj.P<0.05.

